# Percolation of Microparticle Matrix Promotes Cell Migration and Integration while Supporting Native Tissue Architecture

**DOI:** 10.1101/2020.08.10.245589

**Authors:** Jeanne E. Barthold, Brittany M. St. Martin, Shankar Lalitha Sridhar, Franck Vernerey, Stephanie Ellyse Schneider, Alexis Wacquez, Virginia Ferguson, Sarah Calve, Corey P. Neu

## Abstract

Cells embedded in the extracellular matrix of tissues play a critical role in maintaining homeostasis while promoting integration and regeneration following damage or disease. Emerging engineered biomaterials utilize decellularized extracellular matrix as a tissue-specific support structure; however, many dense, structured biomaterials unfortunately demonstrate limited formability, fail to promote cell migration, and result in limited tissue repair. Here, we developed a reinforced composite material of densely packed acellular extracellular matrix microparticles in a hydrogel, termed *tissue clay*, that can be molded and crosslinked to mimic native tissue architecture. We utilized hyaluronic acid-based hydrogels, amorphously packed with acellular articular cartilage tissue particulated to ~125-250 microns in diameter and defined a percolation threshold of 0.57 (v/v) beyond which the compressive modulus exceeded 300kPa. Remarkably, primary chondrocytes recellularized particles within 48 hours, a process driven by chemotaxis, exhibited distributed cellularity in large engineered composites, and expressed genes consistent with native cartilage repair. We additionally demonstrated broad utility of tissue clays through recellularization and persistence of muscle, skin, and cartilage composites in a subcutaneous *in vivo* mouse model. Our findings suggest optimal strategies and material architectures to balance concurrent demands for large-scale mechanical properties while also supporting integration of dense musculoskeletal and connective tissues.

## INTRODUCTION

The extracellular matrix (ECM) is a dynamic environment that dictates and is regulated by activity of resident cells within tissue ^1^. The ECM supports macroscale structure and protection for cells under physiological loading, while also providing microscale cellular contact for proliferation, differentiation, and receptor signaling ^2^. Following injury to a tissue, the ECM signals resident cells to upregulate growth factor production and chemical signaling which promotes migration of inflammatory cells to the damaged site to begin a natural regenerative cascade of ECM repair ^2,3^. The specific structure and composition of both healthy and engineered ECM determines its migratory and self-regenerative capabilities.

A critical paradox of regenerative materials, especially for dense tissues, is the need to provide tissue-specific mechanical and biochemical properties while enabling cell migration to promote normal tissue function and repair. From a structural and compositional standpoint, acellular ECM of native tissue is an ideal material for regeneration because it preserves the complex tissue architecture, molecular composition, and structural rigidity. However, the density of native ECM in musculoskeletal tissues can present a barrier to healing because cell migration arrests when the ECM pore size is less than ~10% of the nuclear area ^4,5^ (Figure 1; Supplementary Tables 1 and 2). Moreover, even in dense *in vitro* collagen gels, migration of mesenchymal lineage cells arrest once the collagen density reaches a critical threshold ^6,7^ which is 30-40 times lower than the collagen density in adult connective tissues ^4,8^. Recent work confirms the finding that nuclear stiffness and size are key factors responsible for migration arrest, as mesenchymal cells that were treated to soften the nuclear architecture showed increased migration into dense hydrogels and devitalized connective tissue matrices ^8^. These findings indicate that the regenerative potential in utilizing large implants of dense acellular matrices is limited in part by pore-size constraints ^9^. In contrast, hydrogel-based approaches to tissue repair can provide higher porosity, as 1% HA/PEGDA gels maintain a 10-16 μm pore diameter ^10^, and modifiable biochemical contact sites but often fail to match the mechanical properties of dense connective tissues ^1,11^. The devitalization of a decellularized tissue into small microparticle (i.e., morselized or powderized) fragments has been proposed as a hybrid method to increase the likelihood of cell migration into acellular tissue ECM ^12^. Key to this observation is that cells are capable of limited migration into dense matrix ^9^, and the repacking of cellularized microparticles provides a means to attain a formable, high-density engineered material, comprised of individually cellularized microparticles. While initial studies using microparticulated tissue have shown some evidence of the chemotactic role that the ECM may play ^12,13^, it is unknown whether microparticulated tissues could address the need for mechanical and biochemical properties while enabling cell migration.

**Figure 1:**
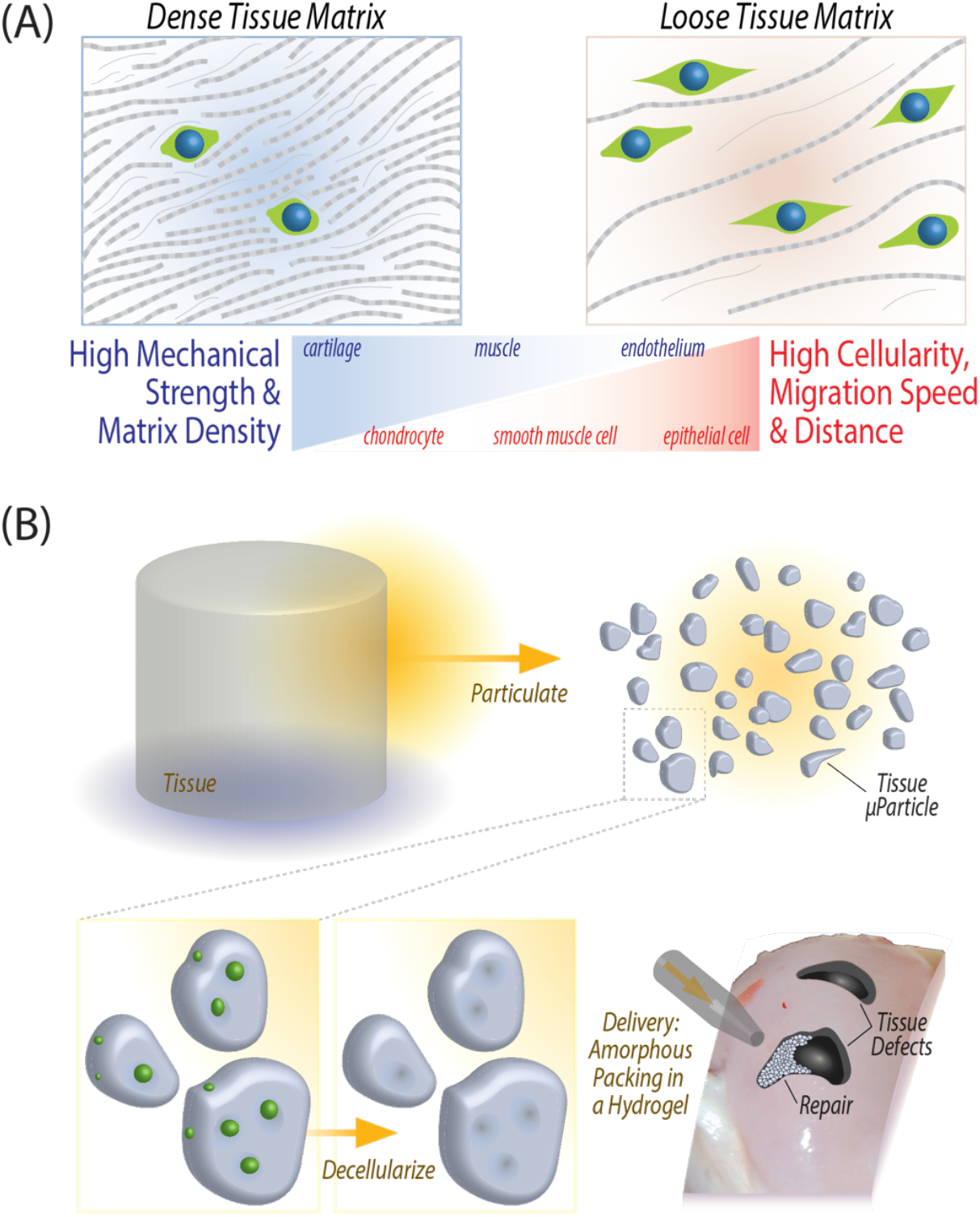
Design of an engineered extracellular matrix based scaffold, *tissue clay*, that maintains high mechanical strength while promoting cell migration and tissue repair. (A) Mechanically robust, dense musculoskeletal tissue withstands high mechanical loads, but the dense ECM also inhibits cell migrate through the tissue for repair after injury. Pore size constraints and nuclear size directly determine the rate of migration to an injury site (Supplementary Tables 1 and 2). (B) Fabrication of *tissue clay* requires pulverization of articular cartilage, decellularization, and amorphous packing in a chondrogenic hydrogel to enable molding in a wide range of defect shapes or sizes.

In this work, we address the paradox of engineering materials of acellular ECM with high mechanical properties and cell migration by creating a *tissue clay*, defined as a reinforced composite material of densely packed acellular ECM microparticles in a hydrogel. Our objective was to develop a general approach for producing tissue clays, which have the benefit of forming and molding to mimic native tissue architecture, packing to high-density to mimic tissue mechanical properties, and enabling cellular migration and recellularization in large defects. Here, we specifically use articular cartilage tissue as a challenging model system: structural architecture is essential to the functionality of cartilage, evidenced by routine loading of multiple times body weight ^14^. Additionally, the density of native cartilage ECM inhibits natural chondrocyte migration and repair of defects ^15^ (Figure 1). Due to the dense ECM structure and mechanical function, articular cartilage presents an extreme test case for our tissue repair method, which we use to test our hypothesis that a decellularized microparticulate ECM hydrogel can enable cellular migration while maintaining native mechanical properties.

## RESULTS

### Tissue clay – a reinforced composite hydrogel of densely packed acellular ECM particles – exhibits structural and mechanical properties approaching that of native tissue

We created *tissue clay* as a combination of a hydrogel material encapsulating acellular ECM particles. Hyaluronic acid (HA) was selected as a promising hydrogel material for cartilage regeneration ^16^ as it is biocompatible and has been shown as a scaffold material that can facilitate nutrient diffusion. Furthermore, hyaluronic acid is an important component of connective tissues and facilitates lubrication, cell differentiation, and cell growth ^17^. When hyaluronic acid is functionalized with thiol groups and combined with polyethylene glycol diacrylate (PEGDA) monomers, the acrylate groups covalently link with thiols through Michael addition at physiological temperatures to create a stable, crosslinked, 3D hydrogel ^10^. As a result, this hydrogel platform can fill a wide range of defect shapes and forms into a stable hydrogel once applied to the injury site. We hypothesized that HA-PEGDA hydrogels would serve as the ideal hydrogel base to encapsulate acellular tissue microparticles.

*Tissue clay* is a biomaterial tailored to mimic native articular cartilage in protein composition and in mechanical properties using the HA/PEGDA hydrogel in combination with acellular cartilage ECM microparticles. To fabricate cartilage *tissue clay*, we decellularized and pulverized native porcine articular cartilage into microparticles smaller than 250 μm in diameter, and then packed the microparticles within the hydrogel at various concentrations (Figure 2). The microparticle concentration in HA/PEGDA hydrogels first increased from a volume fraction of 0 to 0.5, leading to a 5-fold increase in the compressive modulus (~10 kPa to ~50 kPa). To achieve a compressive modulus similar to native tissue, we employed additional centrifugation during hydrogel polymerization, resulting in a packed hydrogel with a volume fraction of 0.6. The increase in density of amorphously packed microparticles from a volume fraction of 0.5 to 0.6 led to a non-linear additional 5-fold increase in the compressive modulus, rapidly reaching approximately 300 kPa (Figure 2d).

**Figure 2.**
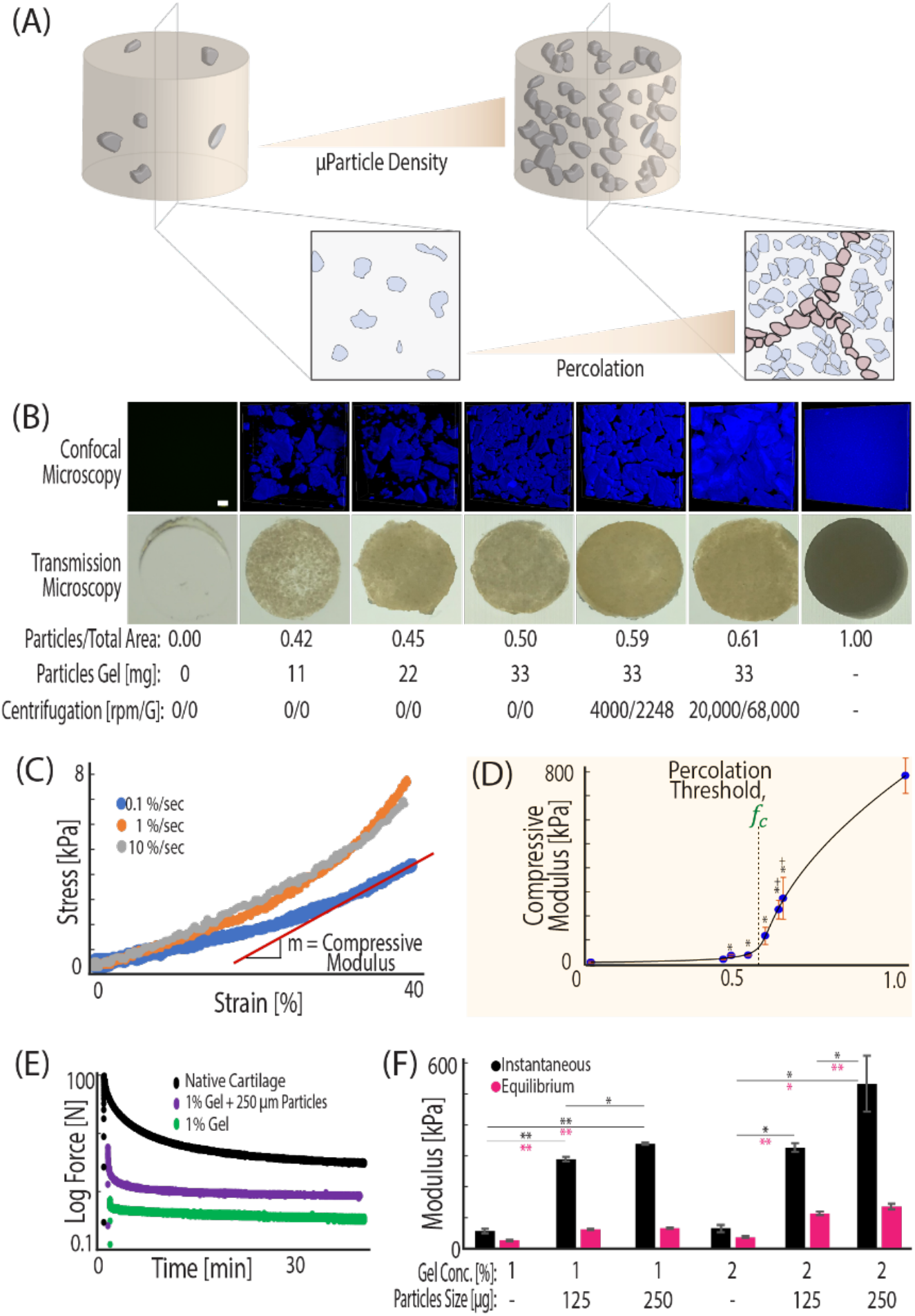
ECM microparticles amorphously packed in a hydrogel beyond a percolation threshold result in a composite material with mechanical properties that mimic native tissue. (A) Packing microparticles to high density in hydrogels increases cell-ECM contact and the volume ratio of microparticle to gel. As the volume ratio increases, the amount of void space between particles decreases until microparticles contact each other and form a new network. (B) The increase in microparticle volume ratio is visualized by taking 405 nm confocal microscopy *z*-stacks of DAPI stained *tissue clay* constructs. (C) Hydrogels were mechanically tested at 0.1%/sec to a deformation magnitude of 40%, and compressive modulus was calculated from the linear portion of the stress vs. strain curve. (D) The modulus of the increasing density *tissue clays* demonstrates an inflection point, *f_c_*. The General Effective Medium (GEM) percolation model fit to the compressive modulus vs. density curve defines the percolation threshold at 0.57 volume ratio: the mathematical point where microparticles are predicted to interact and form a new network to propagate loading. (E, F) *Tissue clay* can be composed of a variety of hyaluronic acid concentrations and microparticle sizes, but for work presented here we utilized 1% HA hydrogels and 250 μm sized particles due to a balance of material consistency and strong mechanics. (*p<.05, **p<.001, +p<.05 as compared with hydrogel before centrifugation).

We hypothesized that the dramatic increase in compressive modulus between 0.5 and 0.6 volume fraction was attributed to packing the microparticles beyond a percolation threshold. Percolation is defined as the point at which the mechanics of the composite system becomes dictated by the network of interconnected stiff constituent pieces, rather than the soft surrounding hydrogel matrix ^18^ (Figure 2). To test our hypothesis, we employed percolation theory that applies across disciplines and explains common natural phenomena involving packing of multiphase materials ^19^. Originally developed to model the mechanical behavior of identical objects suspended in a medium ^20^, percolation theory encounters a common difficulty in biological applications, due to the lack of shape uniformity among materials. A recent adaptation of previous models, the General Effective Medium theory (GEM), bridges percolation and homogenization theories to model continuum mechanics of random multiphase materials ^18^.

We applied the GEM percolation model to explain the mechanical basis for the large increase in compressive moduli between *tissue clay* with volume fractions of 0.5 and 0.6. The model considers the individual moduli of both the hydrogel and the cartilage and applies scaling factors to each component (Supplemental Figure 1). As the concentration of microparticles in the hydrogel increases, the microparticles must pack more tightly together. The percolation model supported our hypothesis and demonstrated that increased amorphous packing of microparticles led to a new network of direct microparticle contact (Figure 2a) and resulted in a *tissue clay* that approaches the compressive modulus of native cartilage. According to this model, the percolation threshold for cartilage *tissue clay* lies at a volume fraction of 0.57, and compression past this point led to rapid increases in compressive moduli (Figure 2d).

### Chondrocytes recellularize tissue particles and maintain a chondrogenic expression

We next investigated the fate of chondrocytes when introduced to *tissue clay* with embedded cartilage ECM microparticles compressed beyond the percolation threshold. Surprisingly, when we introduced chondrocytes into the hydrogel portion of the composite, the cells migrated into and recellularized the cartilage microparticles (Figure 3a,b). Time-course imaging determined that cells migrate into the particles within the first two days *in vitro* (Figure 3c), although in some cases we observed recellularization as early as 12 hours after cellular encapsulation in the hydrogel (Figure 3d). Gene expression analysis of the embedded chondrocytes’ RNA indicate that cells in *tissue clay* begin to upregulate key genes such as *SOX9* (a pivotal transcription factor in chondrocytes critical for cell specification and differentiation), *COL2A1* (the predominant collagen type in hyaline cartilage), and *PRG4* (an important molecule involved in boundary lubrication), and significantly downregulate key fibrocartilaginous genes such as *COL1A2*, as compared to chondrocytes plated on tissue culture plastic (Figure 3f). These data suggest not only that chondrocytes can migrate within the decellularized microparticles, but also that this method of artificial tissue production creates a favorable environment to facilitate a chondrogenic lineage and phenotype.

**Figure 3:**
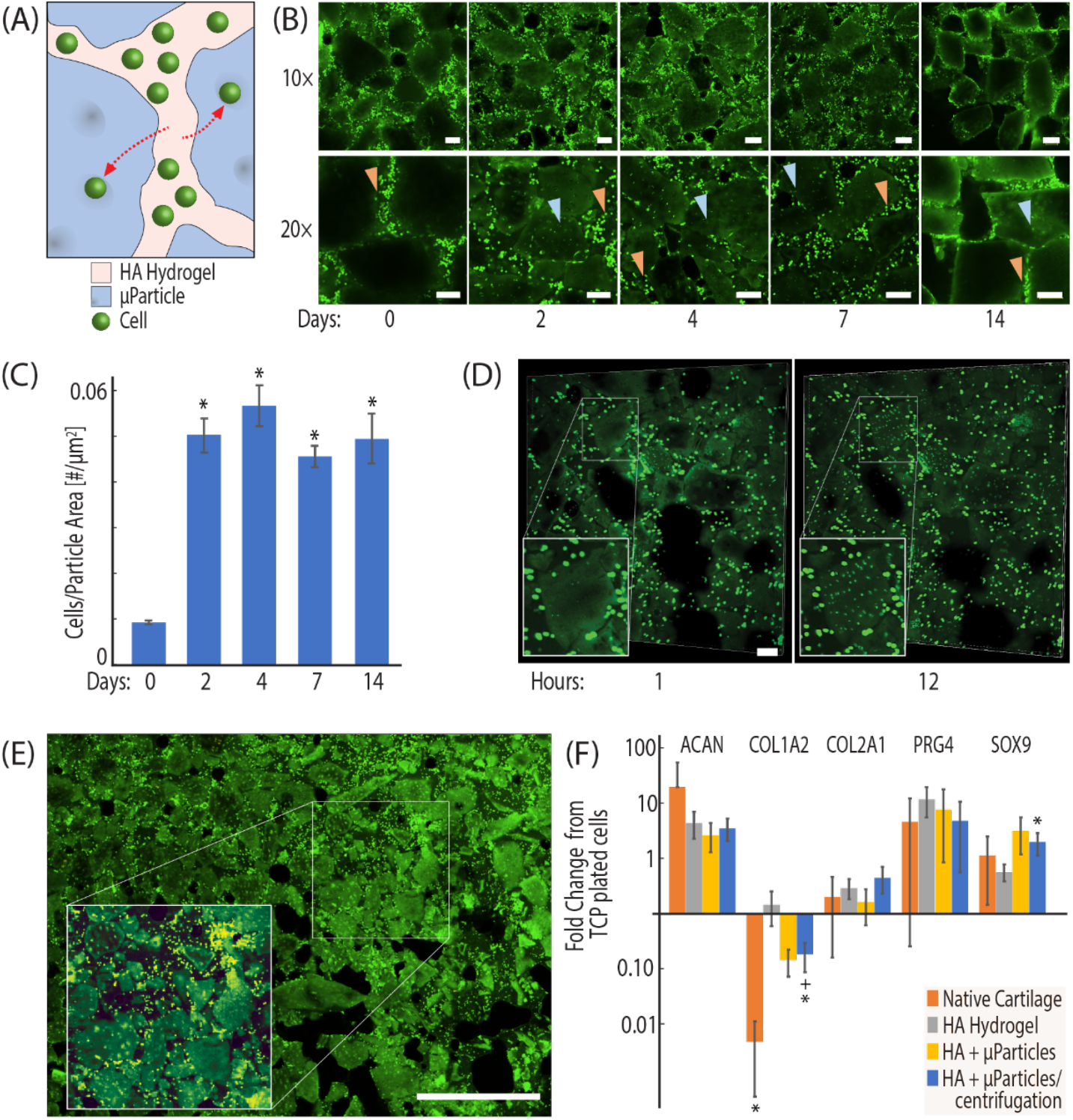
Chondrocytes introduced to the hydrogel region of the composite recellularize microparticles and maintain chondrogenic expression. (A, B) Chondrocytes are extracted from bovine knee joints, stained with a fluorescent proliferation dye (CFSE), and encapsulated in the hydrogel portion of *tissue clay*. The number of cells migrating into particles was measured by quantification of CFSE stained cells present within image-thresholded microparticles at 0,2,4,7, and 14 days post seeding. (C) Chondrocyte migration into the microparticles occurs within the first 2 days of culture. (D) Cells were observed migrating into the microparticles within the first 24 hours. (E) Image montaging supports that the migration was observed broadly throughout the *tissue clay* construct. (F) Quantitative RT-PCR of primary chondrocytes seeded in empty constructs, constructs at percolation, and constructs past percolation shows increased chondrogenic expression in *tissue clay* (HPRT1 used as housekeeping gene, fold change from TCP plated cells). Scale bar = 100 μm (B, D), 1 mm (E). (*p<0.05, RT-qPCR data compared to hydrogel control and ^+^p<0.05, compared to native tissue).

### Chondrocyte migration into tissue particles is attributed to chemotaxis

Considering chondrocytes show limited migration *in vivo* in healthy tissue, and that dense cartilage ECM usually restricts cell mobility ^21,22^ (Figure 1a), the reorganization and migration of chondrocytes into the microparticles was unexpected. We hypothesized that chondrocyte migration into the void regions of acellular cartilage microparticles could be attributed to growth factor reservoirs preserved through decellularization. While sodium dodecyl sulfate (SDS) is typically considered a relatively harsh chemical detergent that could disrupt or wash away growth factors during decellularization ^23,24^, we found no significant reduction of one primary growth factor, TGF-β1, in decellularized particles as compared to native tissue (Figure 4). Additionally, using a standard transwell migration assay, we investigated whether decellularized cartilage microparticles induce nearby cell migration with chondrocytes that are not encapsulated in our hydrogel system (Figure 4a) and found the migration gradient of the microparticles is significantly stronger (p<0.05) than that of standard chondrocyte media (DMEM-F12 with 10% FBS) and nonsupplemented DMEM (Figure 4b). Growth factor reservoirs within acellular ECM microparticles influence the migratory behavior of primary chondrocytes, partially explaining the chondrocyte migration into microparticles that we observed within the first two days of culture.

**Figure 4:**
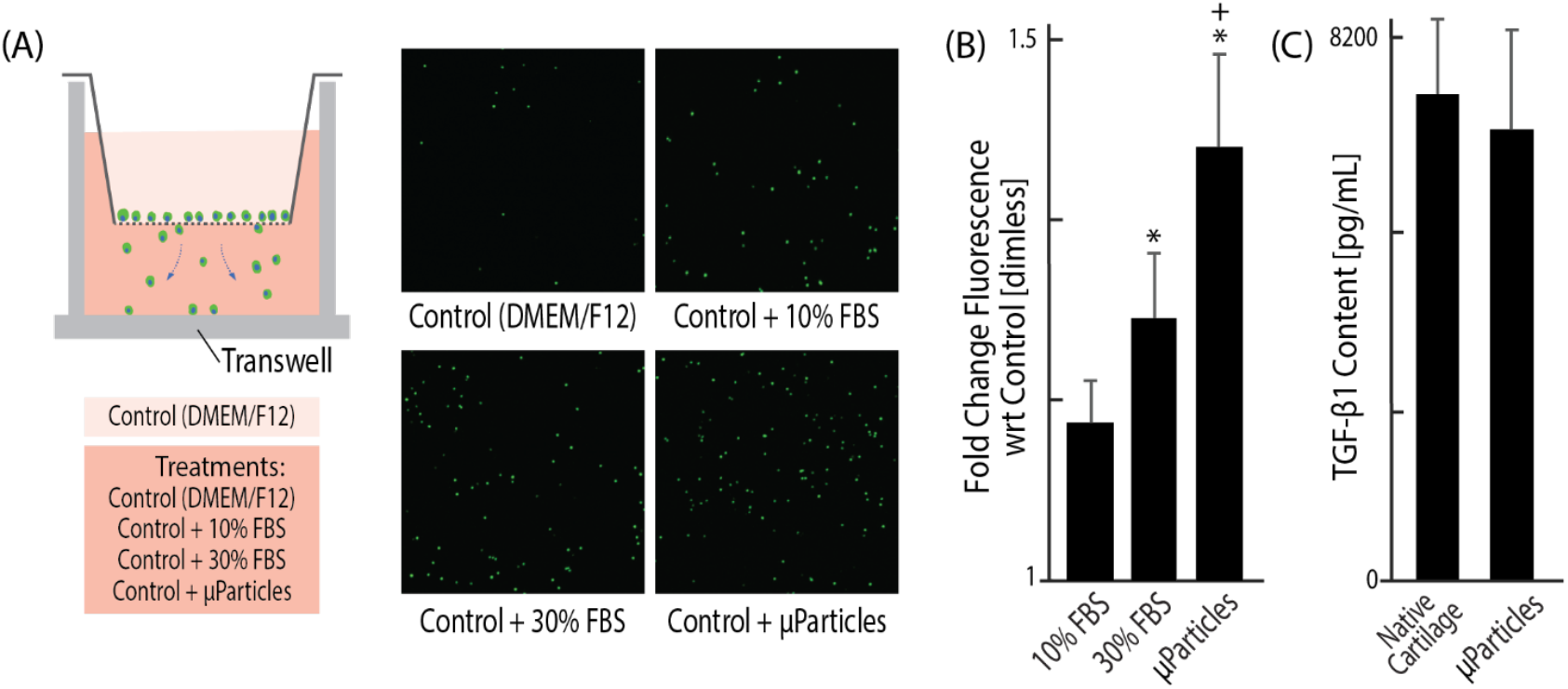
Growth factors are conserved in an acellular extracellular matrix and create a chemotactic gradient which promotes chondrocyte migration into tissue particles. (A, B) Using an established transwell migration assay with four different chemotactic gradients, typically non-migratory chondrocytes demonstrate a significantly greater attraction to the acellular ECM particles than to typical chondrocyte DMEM-F12 media, even with 10% FBS. (C) TGF-β is shown to be highly conserved in the microparticles, despite the processes of decellularization, which likely contributes to the chemotactic influence of the microparticles. (p < 0.05, *compared to DMEM, ^+^compared to 10% FBS).

*Tissue clay* overcomes the common imbalance of engineered materials that are optimized for either strong mechanics or cellularity as *tissue clay* retains mechanical properties of dense native tissue while also enabling cellular migration. Recently published research shows a few key factors ^21^ dictate cellular migration, including chemotactic signaling and material porosity. Migration in our system is likely affected by both factors in tandem, but the chemotactic effect of remaining growth factors plays a distinct role (Figure 4). Additionally, we know that the porosity of the HA/PEGDA hydrogel is dictated by the percentage of thiolated groups on the HA molecules, as well as by the ratio of thiol groups to diacrylate groups. We chose to use a 1% cross linked hydrogel, rather than a 2% hydrogel, to maintain larger pores (~15 μm pore diameter ^10^) for facilitating migration within the hydrogel domains of the *tissue clay* ^25^. Furthermore, chondrocyte migration into tissue ECM has been previously observed at the edges of large acellular ECM implants, but only extending into the tissue about 25-100 μm from the edge ^26–28^. Thus, pulverizing the cartilage ECM until the microparticle diameter is <250 μm and embedding in a hydrogel with 15 μm pores facilitates chondrocyte migration towards matrix-bound TGF-β1. Finally, acellular particles contain some zones of ~10-15 μm empty pores in regions of previously formed chondrons (Supplementary Figure 2). In follow up studies, tissue particles should be extensively characterized to confirm porosity changes that enable the cells to migrate towards a chemotactic signal in the matrix.

### Tissue Clay is a broadly applicable platform design for regenerative hydrogels that promotes in vivo cellular infiltration without sacrificing mechanical properties

Toward broader applications, our *tissue clay* materials provide a universal and commercially available support material, and by testing different porcine tissue types and packing densities, we demonstrate that our hydrogels support cartilage, muscle, and skin regeneration and are viable in a subcutaneous mouse model (Figure 5). Furthermore, the *tissue clay* is mechanically tunable because the packing density of particles directly influences compressive modulus. We created hydrogels of three unique tissue types by encapsulating size-sorted, decellularized microparticles from the three distinct tissues into the HA/PEGDA hydrogel (Figure 5a).

**Figure 5:**
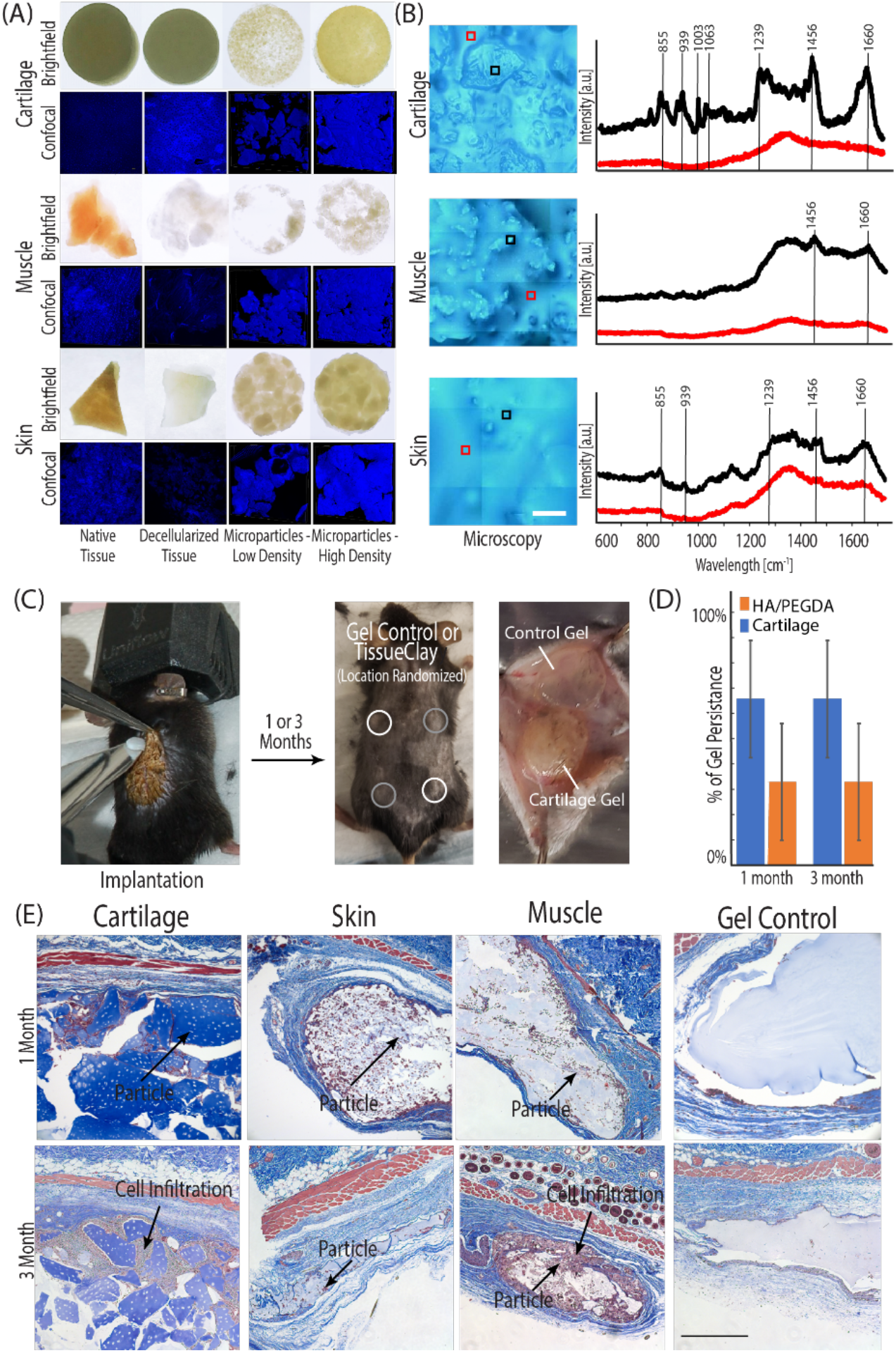
The *tissue clay* fabrication technique is versatile and mechanically tunable for potential successful application with a variety of tissues. (A, C) *Tissue clay* formed from acellular cartilage, skin, and muscle microparticles) were implanted subcutaneously into 8-week old male B6 mice. (B) Prior to implantation, the Raman spectra (representative spectra displayed in B) for all decellularized tissues (e.g. porcine skin, muscle, and cartilage illustrated in black) and surrounding HA/PEGDA hydrogel (illustrated in red) confirmed unique structure signatures in each acellular tissue particle and displayed peaks typical of the respective native tissue, when compared to literature. Each spectra displayed in panel (B) is an average of 9 individual spectra collected in a 30 μm^2^ grid. During the implantation procedure, each mouse received four hydrogels: two HA/PEGDA controls and two *tissue clay* implants. (D) After 1 or 3 months, constructs were explanted and analyzed for native cell infiltration and new ECM deposition. The presence of cartilage tissue microparticles increased longevity of hydrogels in an initial long-term persistence study measured by the presence or absence of the hydrogels at 1 and 3 months. (E) Cell infiltration and new ECM deposition (dark purple nuclei, blue collagen) is observed between and inside of dECM particles towards the edges of the constructs, increasing towards the center of the implant by 3-months (E). Scale bars: B=100 μm, E 500 μm).

Given that our method involves decellularizing tissue to produce microparticles, we first used Raman spectroscopy, and compared our results to known native tissue protein signatures published in literature, to determine whether unique protein macrostructures were maintained through the process of decellularization in each respective tissue. In all tissue types, the spectra collected from tissue particles were distinct from the surrounding HA/PEGDA hydrogel. Additionally, key peaks and relative peak height ratios in our acellular tissue particles compare well with spectra reported previously in native cartilage, muscle, and skin ^29–31^, and the elevated signal intensity, particularly at higher wavenumbers, indicates that collagen and GAG structures native to each tissue type were retained within our decellularized tissue. These results further suggest that the chemistry and structures needed to maintain the biochemical and mechanical characteristics of each tissue persist despite decellularization.

We investigated the translational potential of the *tissue clay* platform via a subcutaneous mouse model. We implanted hydrogels composed of decellularized and microparticulated skin, muscle, or cartilage into male B6 mice (Figure 5c). In the control HA/PEGDA hydrogel, the biomaterial induced a common adverse immune reaction of fibrotic encapsulation: formation of a fibrotic cell capsule directly surrounding the biomaterial to prevent further interplay between biomaterial and host ^32^. By contrast, in the *tissue clay* implants, we observed extensive host cell infiltration into the introduced composite material, limited fibrotic encapsulation in articular cartilage hydrogels, and localization of the cells both around and inside of the tissue particles (Figure 5e). Explants at three months exhibited increased cell infiltration into the *tissue clay* and increased levels of ECM deposition compared to explants at one month (Figure 5e). The particles provide attachment sites, growth factors, and biochemical signals to promote production of extracellular matrix and host integration with the hydrogel. While further work is needed to classify whether cell integration *in vivo* was a macrophage response or a remodeling response, the hydrogels showed increased infiltration over the three months, a time scale longer than a typical initial immune response, suggesting cell homing and matrix deposition, rather than degradation (Figure 5d). All mice survived until sacrifice, with no systemic adverse reactions or pain associated with the implants, confirming the safety of the *tissue clay* materials.

## DISCUSSION

A significant challenge for the design of biomaterials involves providing tissue-specific mechanical and biochemical properties while enabling cell migration to promote normal tissue function and repair. In this work, we meet the dual challenge of structure and cellularity by engineering a native tissue-based hydrogel with particulated ECM to facilitate cellular migration. The unique properties of *tissue clay* are achieved by packing acellular ECM microparticles past the percolation threshold, thereby increasing the compressive modulus and ensuring cell-microparticle interaction, while not inhibiting cellular mobility. We thus developed a material that makes cellular encapsulation simple and provides attachment sites to promote the tissue-specific signaling pathways necessary for growth and regeneration. Cellular migration is especially restricted in dense articular cartilage tissue after injury because the pore size of the ECM (only a few nanometers) is far smaller than the nuclear diameters of chondrocytes and stem cells (~5 um) that could migrate from the bone marrow into the tissue after damage ^33,34^. Utilizing this extreme example, we demonstrate that chondrocytes introduced to *tissue clay* localized both around and within the microparticles and led to an upregulation of key chondrogenic markers by the encapsulated cells. In previous studies using osteochondral implants as a regenerative biomaterial, the tissue matrix was too dense for cells to migrate into the ECM ^35,36^. The work presented here shows that pulverizing ECM and reconstituting the particles in a chondrogenic hydrogel allows for cellular migration while maintaining mechanical properties of the osteochondral implant. These findings are encouraging for regeneration of articular cartilage in future functional *in vivo* studies with the *tissue clay*.

Acellular extracellular matrix as a regenerative material provides tissue specific cell receptors, growth factors, and protein compositions. Recently, several researchers have used digested particles from acellular tissues—e.g. from dermis, bladder, heart, and adipose—to form disassociated tissue-based liquid hydrogels. These hydrogels show significant cellular infiltration *in vivo*, but are structurally inadequate ^37,38^. While these studies support the notion that acellular matrix promotes cellularity, they do not explore the tradeoff with structural stability. Our simple technique of micronizing and decellularizing cartilage extracellular matrix, then recombining the tissue particles with a chondrogenic support matrix, can additionally be optimized for other tissues and applications by varying the packing density and tissue source of particles. *Tissue clay* utilizes the structure and composition of native extracellular matrices and thus mimics the mechanics of a native environment. At the same time, embedded tissue microparticles facilitate cell signaling, inducing native cells to generate their own tissue specific proteins and basement membrane.

We anticipate that our method will aid in overcoming a major difficulty of many previous tissue engineered solutions: that of integrating the implant with the native surrounding tissue ^39^. *In vivo* subcutaneous mouse implants of all tissue constructs found that the material promotes cell infiltration. In articular cartilage *tissue clay*, we additionally observed increased staining for collagen fibers between the microparticles suggesting that the material also promotes matrix production; a promising platform for cell communication and integration. The studies presented here are limited in showing tissue-specific functional behavior. We have shown that our platform provides tissue specific protein structure and tunability to improve mechanics, which together promote cell infiltration, cellular homing, and new ECM deposition. Our platform method enables future work to investigate the *in vivo* functional response in cartilage, skin, and muscle defect models.

## METHODS

### Preparation of Tissue Microparticles

All tissue was sourced from market weight porcine animals within 48 hours of slaughter. Articular cartilage was harvested by exposing the knee joint space and removing tissue with a scalpel, taking care not to include calcified tissue ^9^. Skin was obtained from the hairless stomach region, which we separated from underlying subcutaneous tissue and cut into small pieces prior to storage. Muscle was harvested directly from the thigh of the animal, using a scalpel to cut tissue into small portions. All tissues were frozen at −80°C until further processing. For microparticulation, tissues were pulverized using a liquid nitrogen magnetic freezer mill ^13^, and were sorted via a micro sieve stack to isolate cartilage microparticles <250 μm in diameter, and skin and muscle microparticles <750 μm in diameter (Electron Microscopy Sciences, Hatfield PA). Microparticles were decellularized in 2% SDS for 8 (cartilage), 30 (skin), or 24 (muscle) hours at 37°C, and in 0.1% DNase for 3 hours to remove cellular and genetic material ^40^. Microparticles were rinsed in PBS for 2 hours under agitation, flash frozen in liquid nitrogen, and lyophilized. *Formation of Composite Hydrogels*

Tissue microparticles were encapsulated in hyaluronic acid (HA) hydrogels at high density, facilitating amorphous packing. Lyophilized HA that had previously been thiolated to 25%, which dissolved easily when introduced to sterile DPBS (Hyclone), and formed hydrogels using a Poly(ethylene) glycol diacrylate (PEGDA) cross linker (Alfa Aesar) with a ratio of 1:0.8 thiols to PEGDA. Two aqueous solutions (with a final ratio of HA 10 mg/ml and PEGDA 8.6 mg/ml) were combined and then either plated or mixed with microparticles (at varying ratios, maximum ratio was 220mg microparticles/mL HA/PEGDA hydrogel) prior to plating. Both solutions were plated in custom, standardized molds made from PDMS with glass slides on the top and bottom. The control HA/PEGDA hydrogel and microparticle composite hydrogels were incubated at 37°C for 30 min to facilitate Michael addition crosslinking of the diacrylate groups on the PEGDA with the thiol groups on the HA molecules to form a stable 3D structure. In some sample groups, to increase packing density, a PDMS mold was used with the HA/PEGDA plus microparticles solution and was placed in a centrifuge at 4000 rpm for 20 minutes during polymerization. This helped to compact and evenly distribute microparticles throughout the depth of the hydrogel, confirmed with confocal microscopy (Supplementary Figure 3).

### Confocal Imaging and Calculation of Tissue Microparticle Density

Composite hydrogels were washed twice with PBS, followed by staining for 10 minutes with a standard DAPI stain at a concentration of 5μl stain/ 1 ml DPBS. Hydrogels were imaged on an inverted Nikon A1 confocal microscope with a standard 405 nm laser and 10× objective. Using ImageJ software, confocal images were thresholded ^41^ to identify microparticles within the image regions of interest. The thresholding mask was transformed into an outline, where the area within the microparticles was divided by the total image area to define microparticle density within the region of interest (Supplementary Figure 4). We imaged each composite hydrogel at 3 unique locations, and averaged the microparticle:hydrogel area fraction among the three locations. *Mechanical Testing and Calculation of Percolation Threshold*

Mechanical properties of composite hydrogels were assessed using unconfined compression testing on a Bose ElectroForce 5500 system. We compressed the hydrogel to a strain of 40% of the initial height (0.1% per second) to ensure quasi-static loading following initial (0.1 N pre-load) contact (Figure 2c). The percolation threshold was determined using a General Effective Medium model ^18^, which includes the compressive modulus of both the tissue microparticle and hydrogel constituents, and a custom MATLAB power law analysis.

### Cell Isolation

All primary cells were sourced from juvenile bovine knee joints within 12 hours of slaughter. Chondrocytes were harvested by exposing the knee joint space and removing articular cartilage (superficial and middle zones) with a scalpel ^9^. To extract primary chondrocytes, cartilage was rinsed in sterile PBS, diced into <1 mm^2^ pieces, and digested with 0.2% collagenase-P (Roche Pharmaceuticals) added to serum-free and chemically-defined medium, specifically Dulbecco’s Modified Eagle Medium: nutrient mixture F12 (DMEM-F12) supplemented with 0.1% bovine serum albumin, 100 units/mL penicillin, 100 ug/mL streptomycin, and 50 ug/mL ascorbate-2-phosphate.

### Recellularization of Primary Chondrocytes into Microparticles

For all migration studies, freshly extracted chondrocytes were stained with carboxyfluorescein succinimidyl ester (CFSE) (Invitrogen) prior to hydrogel encapsulation. The chondrocyte cell pellet was resuspended in a 5μM CFSE solution, incubated for 20 minutes at 37°C, and quenched using serum-supplemented (10% FBS) defined medium for 5 minutes at 37°C. Fluorescently labeled chondrocytes were suspended in HA/PEGDA hydrogel solution (1×10^6^ cells/ml), and then mixed with lyophilized microparticles to form cell-laden composite hydrogels, or directly formed without microparticles for control hydrogels). After crosslinking, hydrogels were suspended in chondrogenic medium (DMEM-F12 supplemented with 10% FBS, 0.1% bovine serum albumin, 100 units/mL penicillin, 100 ug/mL streptomycin, and 50 ug/mL ascorbate-2-phosphate) for 14 days, with media replacement every other day. Hydrogels with encapsulated chondrocytes were imaged using a 488 nm laser on an inverted Nikon confocal microscope to define chondrocyte movement throughout the 2-week culture period (on days 0, 2, 4, 7, and 14). Recellularization within the microparticles was quantified using a custom thresholding and counting technique (Figure 3), and staining protocols were optimized and confirmed (Supplementary Figures 5 and 6). Each hydrogel was imaged at three independent locations, and the number of cells/particle area was averaged among the three locations.

### Gene Expression of Chondrocyte-laden Hydrogels

To quantify gene expression of cell-laden composite hydrogels, freshly isolated chondrocytes were resuspended in HA/PEGDA solution at a cell density of 1×10^6^ cells/ml prior to hydrogel formation and culture, as described previously. After 14 days, *tissue clay* and control hydrogels were homogenized for 2 minutes (TissueRuptor) in QIAzol lysis Reagant (Qiagen) and chloroform was added to precipitate and remove proteins. RNA was isolated from cultured cells using the E.Z.N.A Total RNA kit (Omega Tek). Total extracted RNA was reverse transcribed into complimentary DNA (cDNA) (Bio-Rad) using a thermocycler. Quantitative Real-Time PCR (CFX96 Touch, Bio-Rad) was performed on the cDNA using Advanced SYBR Green Supermix and the CFX96 Touch thermocycler (Bio-Rad,). For all samples, HPRT1 was utilized as the housekeeping gene. Known genes for chondrocyte differentiation (SOX9, Col1A2, Col2A1, ACAN, and PRG4) were quantified in all groups: HA/PEGDA hydrogel, HA/PEGDA hydrogel with microparticles, HA/PEGDA hydrogel with microparticles and polymerized under centrifugation, and chondrocytes plated on tissue culture plastic. All measurements were normalized to the housekeeping gene, and fold changes were measured from gene expression of tissue culture plastic plated cells. All primers were designed to be specific for all known isotopes and separated by at least on intron or span an exon-exon junction, if splicing information was available.

### Transwell Migration Assay

To quantify cell migration in response to chemical stimuli, CFSE stained cells were plated on the apical surface of Corning Fluoroblok 24-well plate inserts immediately following staining (40,000 cells in 200 μl of standard chondrocyte medium). In parallel, a known chemoattractant, experimental sample, or controls were added to the basal chamber (600 μl): DMEM-F12 with 10% FBS (standard chondrocyte medium), DMEM-F12 with 30% FBS (enhanced serum medium), DMEM-F12 with 10mg/mL homogenized acellular ECM microparticles (experimental sample), or serum-free DMEM-F12 (negative control) (Figure 4a). All Fluoroblok wells were incubated for 12 hours at 37°C, 5% CO2. After 12 hours, inserts were transferred to a new clear bottom plate where fluorescence was immediately measured using a bottom reading fluorescence plate reader (BioTek, Agilent) at 485/535 nm (Ex/Em). All results are presented as a fold change from the negative control (serum-free DMEM-F12). Supporting images were taken of the same Fluoroblok inserts on an inverted Nikon confocal microscope, 10× objective as previously described.

### TGF-β Quantification to Evaluate Effect of Decellularization on Growth Factor Concentration

TGF-β levels were measured in microparticles before and after decellularization using the Quantkine ELISA kit (Bio-Techne, R&D Systems), following manufacturer instructions. Tissue particles were homogenized in serum-free DMEM F-12 and supernatant collected prior to testing. *Raman Spectroscopy to Evaluate Composite Hydrogel Structure*

Raman Spectroscopy was performed using an upright InVia microscope (Reinshaw, Wotton-under-Edge, UK). Regions of interest on each sample were identified using brightfield microscopy at 5× magnification. Hydrogels were submerged in PBS to eliminate drying, and a 63× immersion objective was used in tandem with a 785 nm laser to excite a 1.064 μm diameter spot on the sample surface. The charge-coupled device (CCD) camera collected reflected spectra for wavelengths between 620 nm and 1711 nm. On each sample (acellular cartilage, acellular muscle, acellular skin, and HA-PEGDA hydrogel), three unique spectral maps were collected, each containing nine points covering 30 μm x 30 μm area. In each spectral map, cosmic rays were identified and removed, a linear baseline was subtracted and intensity normalized, and spectra were smoothed to remove noise ^42^. The nine spectral acquisitions that composed each spectral map were averaged and peaks were identified using an automated peak pick function in Reinshaw WiRE software. The Raman peaks identified in cartilage include C-C stretching (817 nm), hydroxyproline (855 nm), C-C collagen backbone (939 nm), phenylalanine (1003 nm), amide III (1239 nm), CH2CH3 confirmation collagen assignment (1456 nm), and amide I (1660 nm), as well as GAG (1060 nm). Muscle and skin exhibit several typical collagen peaks (855, 1003, 1456, and 1660 nm) (Figure 5b). We used these data to compare to known signatures expected in native cartilage, muscle, and skin tissue ^29–31^.

### In Vivo Implantation and Evaluation via Subcutaneous Mouse Surgery

All animals used in this study were 8-week-old C57BL/6J male mice (#000664) from Jackson Laboratories, and acclimated to surroundings for at least one week prior to the experiment. Protocols were performed in accordance with NIH guidelines for animal handling ^43^ and approved by the Institutional Animal Care and Use Committee by the University of Colorado at Boulder. Animals were housed in a temperature-controlled environment with 12-hour light cycles and received water and food ad libitum. Prior to implantation, endotoxin levels were measured in hydrogels using a gel clot endotoxin detection kit, following instructions from the manufacturer (Genscript). On the day of experiment, mice were anesthetized with isoflurane inhalant and the pedal response was used to test for sensation. Mice were then placed on a warming pad, ophthalmic ointment was applied to the corneas, and nails were trimmed. The surgical site on the mouse was shaved and cleaned with alcohol and betadine solution. Finally, a slow release analgesic (meloxicam) injection was delivered subcutaneously in the peritoneum. Two 8 mm incisions were made in each mouse, one between the shoulders and one between the hips. To the left and right of each incision, a 0.5 cm2 pocket was formed to disrupt the fascia. One hydrogel (microparticle or control), 6 mm diameter and 2 mm thickness, was implanted per pocket, with randomization, and incisions were closed using wound clips. Wound clips were removed once the incision had healed completely. For the first 14 days post-surgery, mice were checked daily for health and signs of infection or pain (swelling, redness of wound, weight loss, isolation, and/or loss of appetite). At 1- or 3-months post-surgery, the animals were euthanized (Supplementary Figure 7). Hydrogels and surrounding tissue were then explanted for analysis.

### Histology of Explanted Hydrogels

Explants harvested from mice after euthanization were fixed in 4% PFA for 48 hours, dehydrated with a sequence of increasing concentration of ethanol (from 70% to 100%), embedded in paraffin, and sectioned using a microtome. Slices were mounted onto microscopy slides and stained with Masson’s Trichome Kit (Newcomer Supply) to analyze structure, cell location, and new ECM deposition.

### Statistics

To test our hypotheses that hydrogel treatment (microparticle density) is able to increase compressive modulus, chondrogenic gene expression, cellular migration, and hydrogel persistence *in vivo*, we performed a separate two-way analysis of variance (ANOVA) for each response variable with treatment and either animal, day, or implant location as fixed main effects. In the chemotaxis (Transwell migration) assay, we used a mixed-model ANOVA with fluorescence as the response variable, with treatment and time as fixed main effects, and animal as a random effect. Post-hoc analyses were performed Tukey’s Honest Significant Difference Test with statistical significance defined as p<0.05. All statistical testing was performed using R software.

## ACKNOWLEDGEMENTS

We are grateful to Kevin Eckstein for providing technical assistance and training for data collection using Raman Spectroscopy. We also thank the histology core at the University of Denver Cancer Center for embedding, sectioning, and staining of explanted mouse tissues. Additionally, we thank Tyler Novak for the cryo SEM images of the acellular microparticles shown in the supplementary material. Finally, we thank Amber Peirce for revision and editing of the manuscript. This work was supported in part by grants to C.P.N.: DoD/CDMRP W81XWH-20-1-0268, NIH R01 AR063712, NSF CMMI CAREER 1349735.

## AUTHOR CONTRIBUTIONS

Conceptualization, J.B. and C.P.N.; Methodology, J.B., B.S.M, S.E.S, S.C., A.W., and C.P.N.; Modeling, S.S. and F.V.; Formal Analysis, J.B.; Animal Approval, S.E.S and C.P.N.; Writing – Original Draft, J.B. Writing – Review & Editing, All authors; Funding Acquisition, C.P.N.; Resources, S.C., V.F. and C.P.N.; Supervision, C.P.N.

## SUPPLEMENTAL INFORMATION

**Supplementary Table 1.** Transwell cellular migration data from select cell types. References are provided at the end of the Supplemental Information.

| Cell Type | Transwell pore (µm) | Nuclear Diameter | 10% Diameter | Pore/10% Nuclear Diameter | % cells migrate | time (hours) | Chemo attractant |
| --- | --- | --- | --- | --- | --- | --- | --- |
| Chondrocyte | 8<sup>1</sup> | 5 | 0.5 | 16.0 | 11%<sup>1</sup> | 24 | PDGF |
| MSC | 8<sup>1</sup> | 6 | 0.6 | 13.3 | 13%<sup>1</sup> | 24 | PDGF |
| Fibrochondrocyte | 3<sup>2</sup> | 9<sup>2</sup> | 0.9 | 3.3 | 0%<sup>3</sup> | 16 | PDGF |
| Fibrochondrocyte | 5<sup>2</sup> | 9<sup>2</sup> | 0.9 | 5.6 | 20%<sup>3</sup> | 16 | PDGF |
| Fibrochondrocyte | 8<sup>2</sup> | 9<sup>2</sup> | 0.9 | 8.9 | 50%<sup>3</sup> | 16 | PDGF |
| Myoblasts | 8<sup>4</sup> | 10<sup>5</sup> | 1 | 8.0 | 10%<sup>4</sup> | 24 | PDGF |
| Epithelial Cell | 10<sup>6</sup> | 5<sup>6</sup> | 0.5 | 20.0 | 83%<sup>6</sup> | 24 | 10% FBS |
| Epithelial Cell | 8<sup>6</sup> | 5<sup>6</sup> | 0.5 | 16.0 | 65%<sup>6</sup> | 24 | 10% FBS |
| Epithelial Cell | 7<sup>6</sup> | 5<sup>6</sup> | 0.5 | 10.0 | 50%<sup>6</sup> | 24 | 10% FBS |
| Neutrophil | 0.45 | 3 | 0.3 | 1.5 | 0%<sup>7</sup> | 20 | IL8 |
| Neutrophil | 3<sup>7</sup> | 3<sup>7</sup> | 0.3 | 10.0 | 10%<sup>7</sup> | 20 | IL8 |

**Supplementary Table 2.** In vivo migration data from select cell types. References are provided at the end of the Supplemental Information.

| Tissue or System | Cell type | Nucleus diameter (μm) | *In Vivo* Pore size (μm) | Pore size/ Nucleus | Pore size/10% Nucleus | Migration rate (μm /min) | Notes on Migration Rate |
| --- | --- | --- | --- | --- | --- | --- | --- |
| **Skin** | Fibroblast | 7<sup>8</sup> | 2<sup>8</sup> | 0.28 | 2.86 | 0.915<sup>9</sup> | 3D collagen |
| **Breast** | mammary epithelial cell | 4.9<sup>10</sup> | 5<sup>6</sup> | 1.02 | 10.23 | 1.5<sup>11</sup> | 2D on collagen |
| **Meniscus** | Fibrochondrocyte | 9<sup>3</sup> | 0.01<sup>12</sup> | 0.001 | 0.01 | 0.0014<sup>2</sup> | Native tissue slice |
| **Muscle** | Smooth muscle cell | 5<sup>13</sup> | 0.44<sup>13</sup> | 0.088 | 0.88 | 0.7<sup>14</sup> | 2D on Polyacryl. |
| **Muscle** | Satellite cells | 4<sup>15</sup> | 0.44<sup>13</sup> | 0.11 | 1.1 | 0.833<sup>15</sup> |  |
| **Cartilage** | Chondrocyte | 5<sup>16</sup> | 0.006<sup>17</sup> | 0.0012 | 0.012 | 0.027<sup>18</sup> |  |
| **Stem Cell** | MSC | 7<sup>19</sup> | 6<sup>20</sup> | 0.86 | 8.57 | 1<sup>21</sup> |  |
| **Bone** | Osteocyte | 2<sup>20</sup> | 0.2<sup>20</sup> | 0.1 | 1 | 0.002 |  |
| **Adipose** | Adipocytes | 5<sup>22</sup> | 10<sup>23</sup> | 2 | 20 | 0.33<sup>22</sup> |  |
| **Immune** | Leukocyte | 6<sup>24</sup> | 0.8<sup>24</sup> | 0.13 | 1.33 | 10<sup>24</sup> | 3D collagen |
| **Immune** | Tcells | 3.1<sup>24</sup> | 0.8<sup>24</sup> | 0.26 | 2.59 | 11<sup>25</sup> | *In vivo* lymph |
| **Cancer** | Breast Sarcoma Cell | 7.7<sup>24</sup> | 10<sup>24</sup> | 1.30 | 13.03 | 0.33<sup>24</sup> | 3D collagen |
| **Immune** | Neutrophil | 2.5<sup>24</sup> | 0.8<sup>24</sup> | 0.32 | 3.17 | 11<sup>24</sup> | 3D collagen |

### Calculation of percolation threshold from Generalized Effective Medium Theory

In this section, we provide the mathematical equation corresponding to the Generalized Effective Medium Theory ^26^ that were used to fit the effective compressive modulus of the composite material.

The effective mechanical properties of composite materials are typically described using the volume fractions and the corresponding mechanical properties of the constituents. In a two-constituent system, the overall effective compressive modulus *E_eff_* is influenced by the spatial distribution and connectivity of the inclusions of each constituent. Multiple theories on percolation of the inclusions have been introduced to model this behavior. The generalized effective medium theory (GEM) has been found to be the most robust as it can provides a general approach to incorporate differences in percolation thresholds arising from the geometry of the inclusions. Let us denote material 1 as the hydrogel with compressive modulus *E*_1_ and material 2 as the native cartilage with compressive modulus *E*_2_ whose pieces are encapsulated in the sample. Let *f* denote the volume fraction of cartilage in the entire sample such that *1* – *f* denotes the volume fraction of hydrogel. The effective compressive modulus *E_eff_* can be determined by solving the following equation

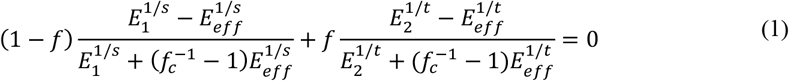

where *f_c_* is the percolation threshold for the volume fraction of cartilage. The terms s and t are the characteristic exponents corresponding to a universal scaling law for effective properties at volume fractions near the percolation threshold *f_c_*. These exponent s captures the sharpness of the transition of the effective modulus from *E*_1_ when *f* ≪ *f_c_* while the exponent t captures the transition to a value of effective modulus of *E*_2_ when *f_c_* < *f* → 1. The figure below shows two plots of the effective modulus *E_eff_* as a function of the volume fraction *f* for different values of *s* and *t*.

**Supplementary Figure 1.**
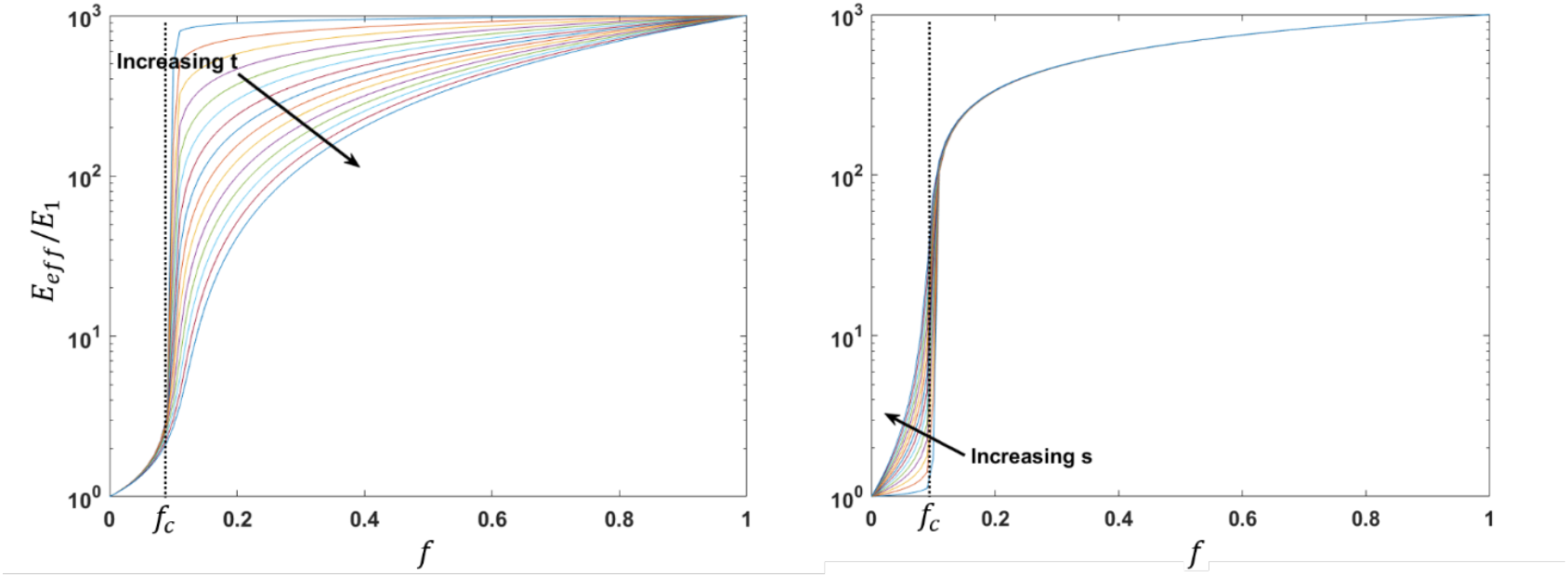
Plots of the ratio of effective compressive modulus *E_eff_*/*E*_1_ predicted by the GEM theory as a function of the volume fraction *f* of cartilage. The difference in the sharpness of transition from the modulus of the hydrogel *E*_1_ to that of cartilage *E*_2_ assumed to be 1000 times higher is illustrated with changes in the exponents *s* and *t*.

**Supplementary Figure 2.**
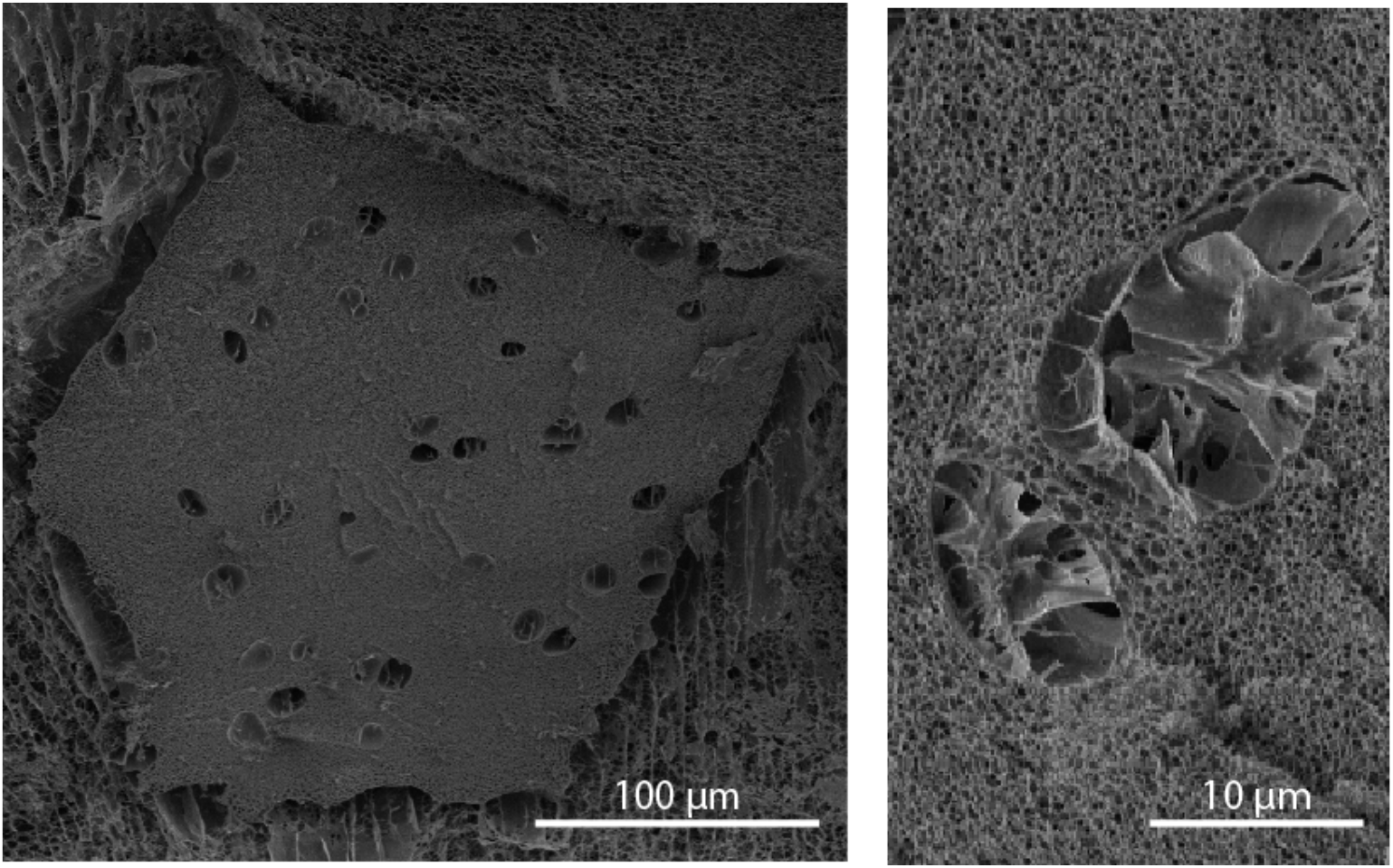
Scanning Electron Microscopy Images of acellular tissue particles. Regions previously formed for chondrons are 10-15 μm in diameter, providing several regions of much larger pores than the surrounding cartilage extracellular matrix.

**Supplementary Figure 3.**
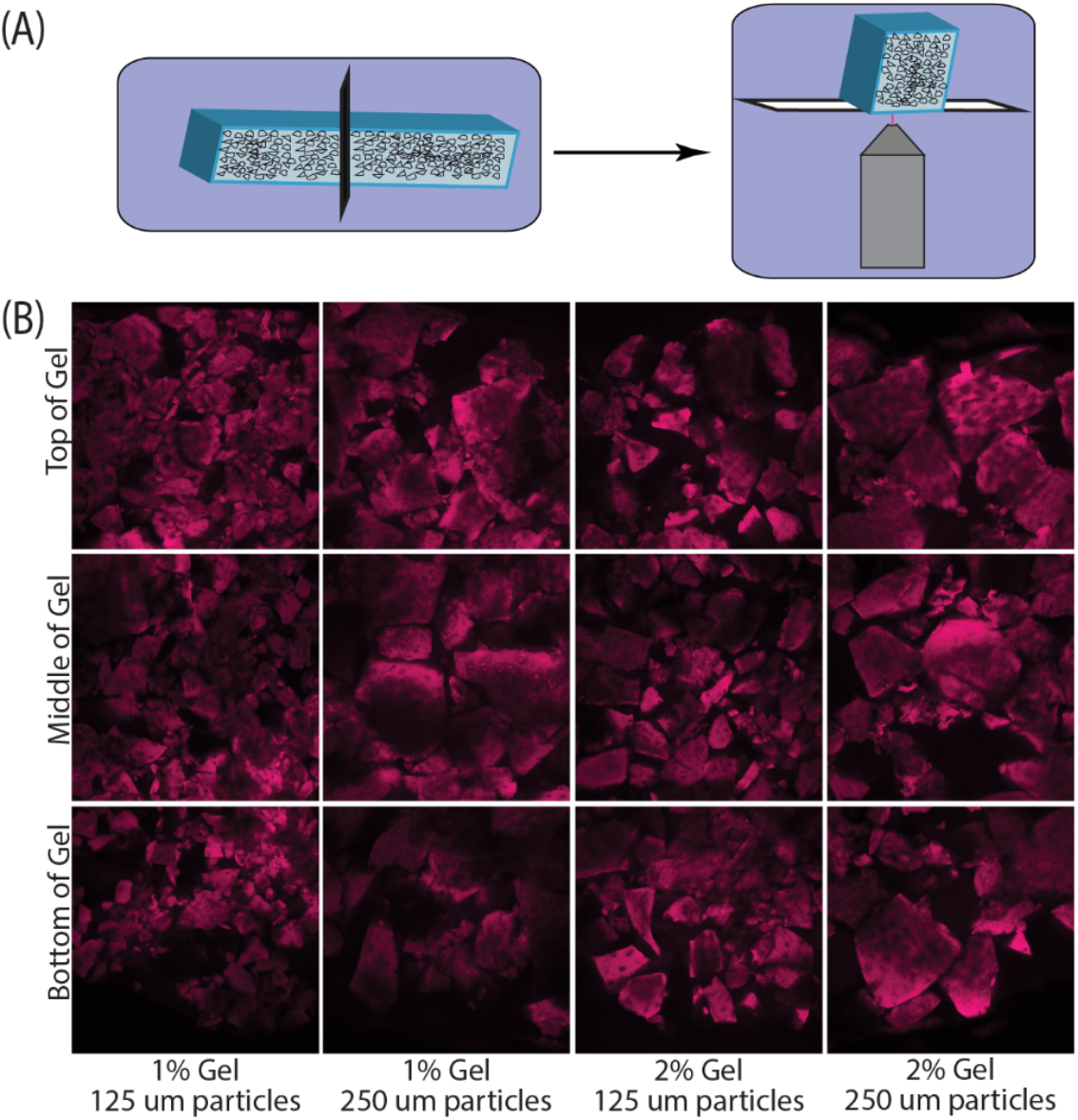
Particle packing was consistent throughout large volumes of the engineered materials. Tissue clays maintain the volume density of particles throughout the depth of the scaffold; particles do not pack in higher density at the base of the scaffold. (A) Polymerized scaffolds are bisected and imaged throughout the depth, (B) Particles of different sizes in HA/PEGDA gels of differing densities pack evenly throughout the depth of the scaffold.

**Supplementary Figure 4.**
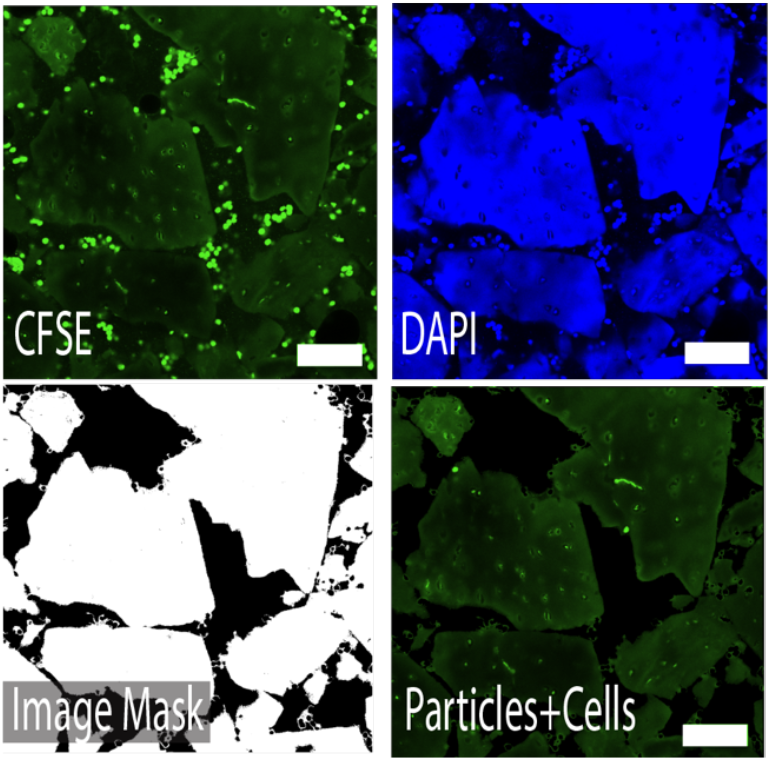
Custom Thresholding enabled quantification of the number of CFSE-stained cells that migrated into particles. Thresholding the particles using DAPI stain in the 405 nm channel allowed us to subtract the background. Once all cells in the hydrogel were subtracted from the background, particle measuring tools in ImageJ enabled quantification of CFSE stained cells in the particles at each time point. Scale Bar = 100μm.

**Supplementary Figure 5.**
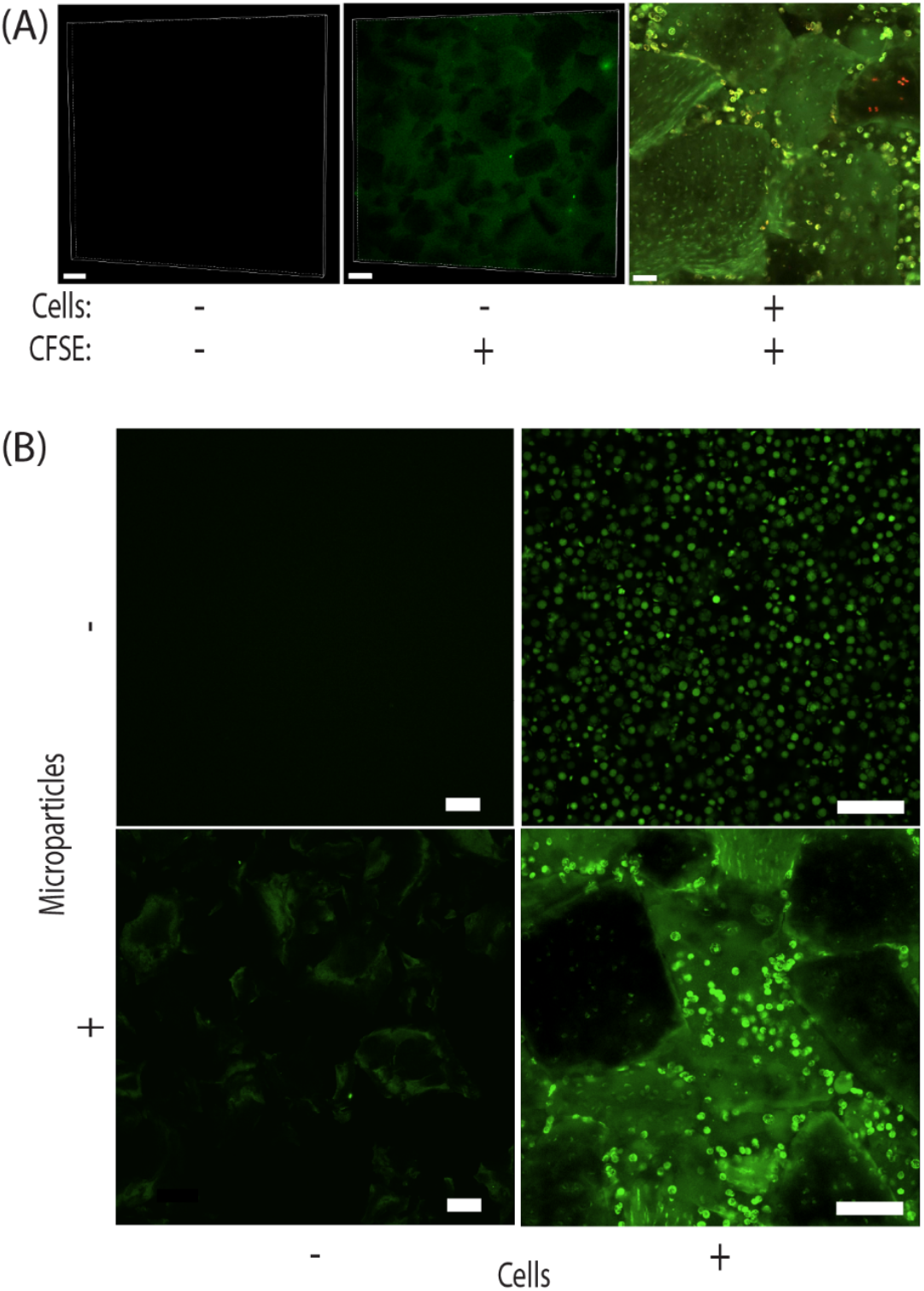
Images confirming CFSE staining of chondrocytes. (A) The addition of representative levels of CFSE to the microparticle gels does not show any bright localization to the particles. (B) Additionally, we do not observe staining due to imaging by confocal microscopy, or the microparticles alone, and we only observe a localization of signal with the addition of cells. Scale Bar = 100μm, final panel of (A), scale bar = 50μm.

**Supplementary Figure 6.**
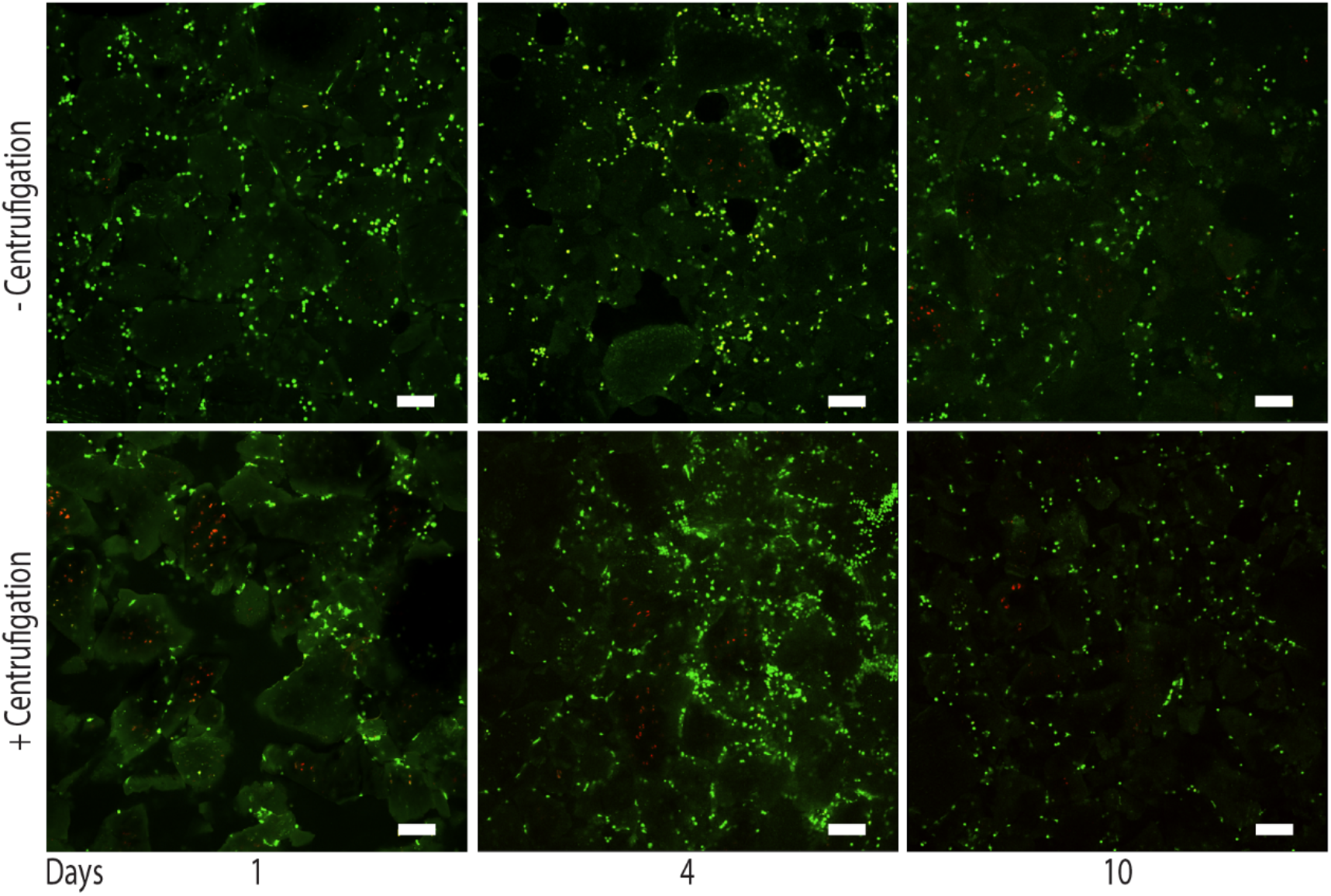
Post-centrifugation live/dead staining. Live dead staining of the cells before and after centrifugation (scaffold packing) demonstrates no dramatic increase in cell death. Green = Live cells, Red = Dead cells. Scale Bar = 100μm

**Supplementary Figure 7.**
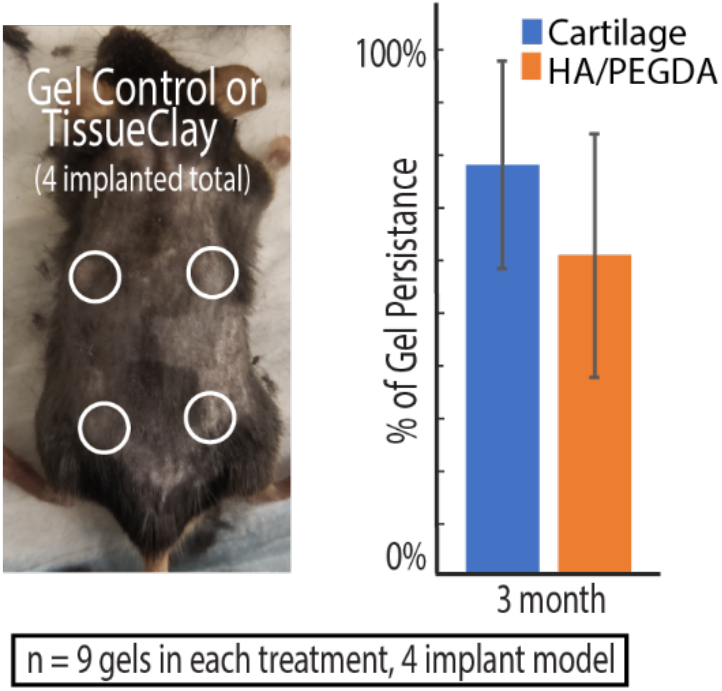
In a supplementary subcutaneous mouse study, we implanted 4 gels (2 *tissue clay* hydrogels and 2 HA/PEGDA controls) in 3 different mice (gel type randomized among the 4 highlighted locations). Data shown here represent the hydrogel persistence over 3 months and the trends agree with the persistence results reported in the main paper.

